# CRISPR interference identifies vulnerable cellular pathways with bactericidal phenotypes in *Mycobacterium tuberculosis*

**DOI:** 10.1101/2021.02.04.429736

**Authors:** Matthew B. McNeil, Laura M Keighley, Josephine R. Cook, Chen-Yi Cheung, Gregory M. Cook

## Abstract

*Mycobacterium tuberculosis* remains a leading cause of death for which new drugs are needed. The identification of drug targets has been advanced by high-throughput and targeted genetic deletion strategies. Each though has limitations including the inability to distinguish between levels of vulnerability, lethality and scalability as a molecular tool. Using mycobacterial CRISPR interference in combination with phenotypic screening we have overcome these individual issues to investigate essentiality, vulnerability and lethality for 96 target genes from a diverse array of cellular pathways, many of which are potential antibiotic targets. Essential genes involved in cell wall synthesis and central cellular functions were equally vulnerable and often had bactericidal consequences. Conversely, essential genes involved in metabolism, oxidative phosphorylation or amino acid synthesis were less vulnerable to inhibition and frequently bacteriostatic. In conclusion, this study provides novel insights into mycobacterial genetics and biology that will help to prioritise potential drug targets.

## Introduction

Infections from *Mycobacterium tuberculosis* remain a leading cause of death^1^. *M. tuberculosis* is currently treated with a combination of four drugs for six months. However, the evolution and spread of drug resistance highlights a need for new drugs. Novel drugs with unique molecular targets that are (i) essential for growth, (ii) are highly vulnerable to inhibition (i.e. require low levels of inhibition to produce a bacterial phenotype)^2-4^ and (iii) are lethal when inhibited are likely to have the greatest impact on reducing treatment times^5^.

Transposon-saturation mutagenesis coupled to next-generation sequencing (i.e. Tn-seq) provides a high-throughput assessment of fitness costs associated with gene disruption. Tn-seq in *M. tuberculosis* has made significant contributions to the identification of essential genes ^6-8^. Despite this, Tn-seq does not distinguish between bactericidal and bacteriostatic outcomes. Although improved statistical analysis has allowed for the identification of differences in genetic requirements between clinical isolates^9^, there is also no standardized Tn-seq methodology to determine which genes are more vulnerable to inhibition^10^. Dual-control (DUC) switches that combine transcriptional repression and proteolysis of target genes overcomes issues associated with Tn-seq^3^. DUC switches can also partially knockdown target genes to generate hypomorphic strains for compound mechanism of action studies and for investigations into pathway vulnerability^2-4^. Despite this, the construction of DUC switches is resource intensive, prohibiting the parallel construction of DUC switches for multiple genes of interest. Notable exceptions include the large collaborative efforts of the PROSPECT screen^11^.

Mycobacterial CRISPR interference (CRISPRi) is an alternative genetic platform that transcriptionally represses target gene expression^12-15^. CRISPRi plasmids are easy to construct as (i) they only require the cloning of only a 20 bp segment of complementarity to target genes (i.e. sgRNA sequences) and (ii) require only a single transformation as CRISPRi plasmids also express the necessary catalytically dead Cas9 (i.e. dCas9) ^13,15^. We and others have shown that CRISPRi can (i) probe gene essentiality, (ii) provide information on the lethality and (iii) can generate partial knockdown strains^12-16^. Using mycobacterial CRISPRi, this current study has phenotypically screened target genes from a diverse spectrum of cellular pathways in *M. tuberculosis* to investigate variations in gene essentiality, vulnerability and lethality. These results demonstrate that the majority of screened genes involved in cell wall synthesis and core cellular functions (e.g. transcription and translation) are essential, equally vulnerable to inhibition and bactericidal when inhibited. Conversely, the majority of essential genes involved in metabolism or amino acid synthesis were subject to buffering effects, requiring higher levels of repression to inhibit growth. Furthermore, the inhibition of several genes previously defined as non-essential inhibited bacterial growth suggesting a delay between target inhibition and the necessary metabolic remodelling to facilitate growth. In conclusion, this study will help to prioritise potential drug targets as well as reiterate the utility of CRISPRi in investigating mycobacterial genetics and physiology.

## Results

### Mycobacterial CRISPRi to assess genetic essentially

Using CRISPRi we investigated the phenotypes (i.e. growth and viability) of 96 genes (encoding 94 proteins and 2 ribosomal RNAs) from a diverse array of biological processes, many of which are considered potential drug targets. Mycobacterial CRISPRi uses a type I dCas9 from *Strepoccous thermophilus* that recognizes 15 permissible non-canonical PAM sequences to repress target gene expression^15^. Both sgRNA and dCas9 are expressed in response to the addition of anhydrotetracycline (ATc). There is a linear relationship between the level of ATc added and the level of transcriptional repression ^13,16^. Two sgRNA sequences per target gene were selected and cloned (Table S1). Some genes have only a single sgRNA due to cloning difficulties. Bacterial phenotypes following target gene repression were screened in 96 well plates using an ATc gradient (Figure 1). OD_600_ was determined 7-days post the addition of ATc and genes were defined as either essential (i.e. <25% growth relative to no ATc control), growth impairing (i.e. <50% growth relative to no ATc control) or non-essential (i.e. >50% growth relative to no ATc control). Essentiality calls were compared with published Tn-eq experiments as available in the mycobrowser database^6^. CFUs were determined 5-days post the addition of ATc as previous work demonstrated that this allows for the detection of bacterial killing prior to the emergence of non-responsive CRISPRi mutants^13,16^. Essential or growth impairing sgRNAs that produced ≥1 log_10_ reduction in CFU/ml were defined as bactericidal, whilst <1 log_10_ reduction was bacteriostatic^13,16^.

**Figure 1:**
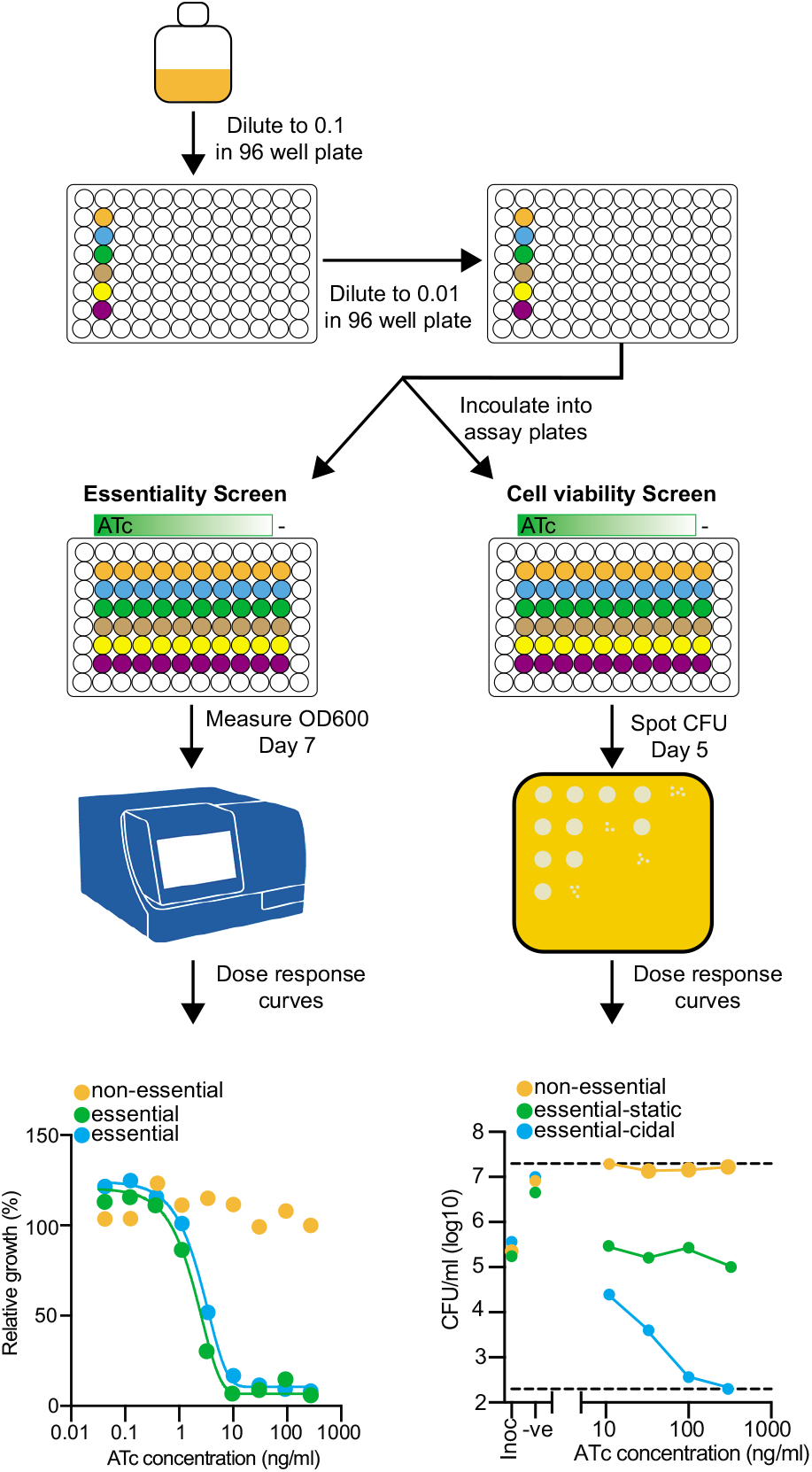
Experimental workflow for CRISPRi assessment of bacterial phenotypes. Cultures of *M. tuberculosis* mc^2^6230 expressing CRISPRi plasmids that express a single sgRNA were maintained in 10 ml cultures. Cultures were diluted to an OD_600_of 0.01 in a deep well 96 well plate and then inoculated into 96 well assay plates that contain a dilution gradient of ATc to induce expression of both the dCas9 and sgRNA at a starting OD_600_ of 0.005. Assays plates used to determine if sgRNAs inhibited bacterial growth were assessed after 7 days by determining the OD_600_relative to a no ATc control. Plates used to determine bacterial viability were assessed after 5 days by spotting for CFUs. Previous work has demonstrated that day 5 allows for the detection of bacterial killing prior to the emergence of non-responsive CRISPRi mutants^13,16^. Further details for experimental workflow are described in the materials and methods

### Cell wall synthesizing genes are essential and bactericidal when inhibited

The majority of sgRNAs targeting genes involved in cell wall synthesis in our screen were essential (Figure 2A). Essentiality and non-essentiality (e.g. *rv3032* and *treS*) calls are consistent with previous experiments (Figure 2A)^6,17,18^. All genes involved in mycolic acid synthesis (i.e. *kasA, inhA, hadA/B, pkS13, fadD32, mmpL3 and pptT*) were essential and bactericidal when inhibited (Figure 2A). The majority of mycolic acid synthesis genes caused a >3 log reduction in CFU/ml at the maximum ATc concentration (i.e. 300 ng/ml) (Figure 2A). Genes involved in arabinogalactan synthesis (i.e. *wecA, embA, dprE1* and *glf*) had either an essential or growth impairing phenotype (Figure 2A). Inhibition of *wecA, embA* and *dprE1* had a weak bactericidal phenotype (i.e. 1 log reduction in CFU/ml) at ATc concentrations >100 ng/ml (Figure 2A). Inhibition of *glf* had a stronger bactericidal phenotype (i.e. >2 log reduction in CFU/ml at lower ATc concentrations) (Figure 2A). All genes in peptidoglycan synthesis (i.e. *murE, murX, murI, alr, ddlA*) were essential for growth (Figure 2A). The inhibition of *murE, murX* and *ddlA* had a bactericidal phenotype, whilst *murI* and *alr* were bacteriostatic (Figure 2A). With the exception of *glgE* that had an essential-acteriostatic phenotype, all genes involved in glucan biosynthesis (i.e. *rv3032* and *treS*) were individually non-essential (Figure 2A). The predicted efflux pump, *efpA*, had an essential-bactericidal phenotype, whilst inhibition of the serine/threonine protein kinase *pknA* had a growth impairing phenotype (Figure 2A). In conclusion, genes involved in cell wall synthesis are predominately essential for growth, although there is variation in whether their inhibition is bactericidal.

**Figure 2:**
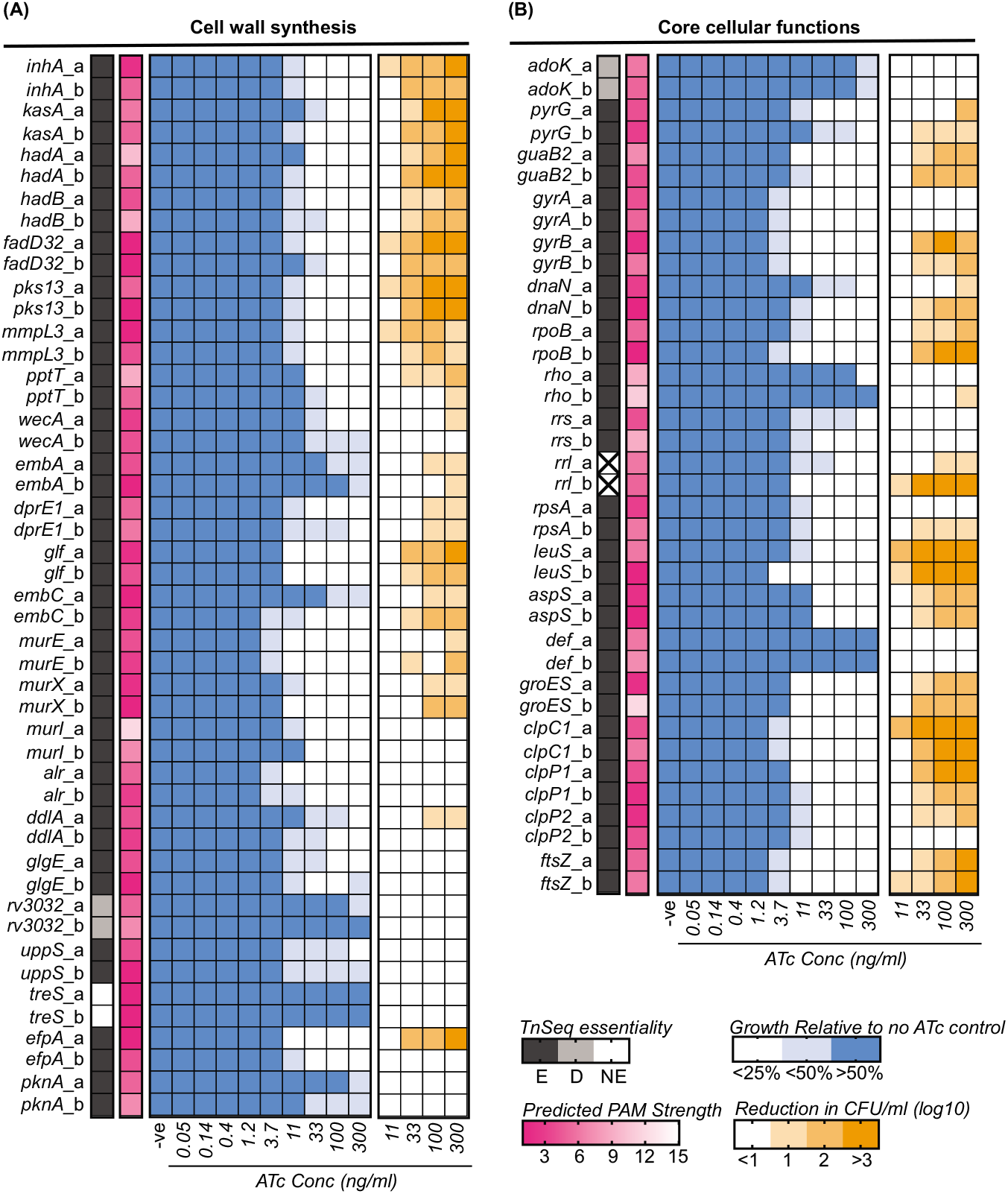
Bacterial phenotypes following the transcriptional inhibition of genes involved in cell wall synthesis or core cellular processes. **(A and B)** Each sgRNA is designated by the target gene name and whether it is the first or second designed sequence (i.e. a or b). Gray shaded heatmap describes the results from previous TnSeq experiments^6^as presented in the mycobrowser database (https://mycobrowser.epfl.ch/) and whether it is essential (E), non-essential (NE) or whether it has a growth defect (D). X denotes genes for which there is no essentiality call. Pink shaded heatmap describes the predicted strength of PAM scores ranked between 1-15 based on previous studies by Rock et al^15^. Blue shaded heatmap describes the level of growth inhibition at increasing ATc concentrations relative to a no ATc control well. The level of inhibition is the mean of at least four replicate experiments. Orange shaded heatmap describes the log_10_ reduction in CFU/ml for each sgRNA between day 0 and day 5 at increasing concentrations of ATc. The log_10_ reduction in CFU/ml of inhibition is the mean of at least four replicate experiments.

### Core cellular processes are essential and have lethal phenotypes when inhibited

The majority of sgRNAs targeting genes involved in core cellular processes (i.e. transcription, translation, nucleotide metabolism, protein homeostasis and cell division) were essential and consistent with previous Tn-seq experiments^6^ (Figure 2B). With the exception of *adoK*, genes involved in nucleotide biosynthesis (i.e. *pyrG* and *guaB2*) were essential-bactericidal (Figure 2B). Both *gyrA* and *gyrB* were individually essential, yet only the inhibition of *gyrB* had a bactericidal phenotype (Figure 2B). Inhibition of DNA polymerase (i.e. *dnaN*) and RNA polymerase (i.e. *rpoB*) were essential and bactericidal, with the phenotypically strongest sgRNAs having a >2 log reduction in CFU/ml. Inhibition of the transcriptional terminator, *rho*, inhibited bacterial growth but only at the highest ATc concentration (Figure 2B). Ribosomal RNAs (i.e. 16S-*rrs* and 23S-*rrl*) were essential for bacterial growth, but only the inhibition of *rrl* resulted in reduced bacterial viability. The ribosomal protein *rpsA* was also essential for growth, with the phenotypically strongest sgRNA reducing bacterial viability. Similarly the transcriptional inhibition of genes encoding tRNA synthetase (i.e. *aspS* and *leuS*), genes involved in protein homeostasis (i.e. *groES, clpC1, clpP1* and *clpP2*) and cell division (i.e. *ftsZ*) were essential and had strong bactericidal phenotypes (Figure 2B). The peptide deformylase encoded by *def* that was previously defined as essential failed to inhibit bacterial growth following transcriptional repression (Figure 2B). In conclusion, genes involved in central cellular processes are essential for the growth of *M. tuberculosis*, and frequently have a bactericidal phenotype.

### Genes involved in metabolic process and oxidative phosphorylation are generally bacteriostatic

Transcriptional inhibition of genes involved metabolic processes or oxidative phosphorylation that had previously been defined as essential resulted in growth inhibition (i.e. *fba, glcB, folP1, folC, atpE* and *atpB*) (Figure 3A and B). However, only the inhibition of *glcB* and *atpE/B* significantly reduced bacterial viability (Figure 3A and B). Some sgRNAs targeting genes previously defined as essential failed to inhibit bacterial growth (e.g. *fum, menA, menD* and *birA*), whilst only the strongest sgRNA targeting *trxB2* was able impair bacterial growth (Figure 3A and B). Furthermore, some sgRNA targeting genes previously defined as non-essential inhibited bacterial growth in our study (i.e. *eno, dlaT, mqo, mdh, icl1, pckA, ndh, qcrB, ctaC* and *ethA*). Whilst the majority were bacteriostatic the strongest *ndh* sgRNA was bactericidal (Figure 3A and B). In conclusion, the transcriptional inhibition of many genes involved in metabolic processes or oxidative phosphorylation inhibit or impair bacterial growth, but have a largely bacteriostatic phenotype.

**Figure 3:**
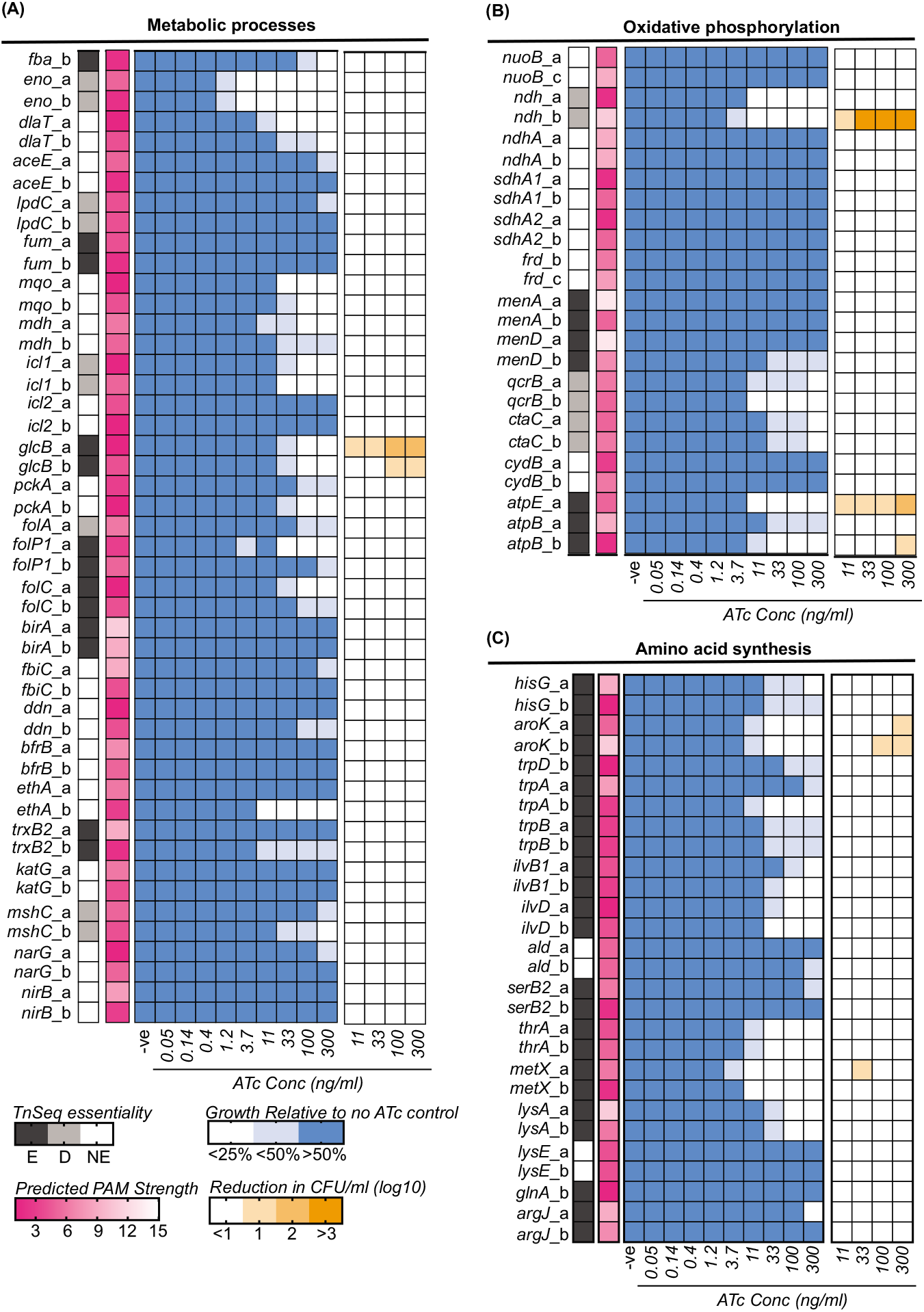
Bacterial phenotypes following the transcriptional inhibition of genes involved in metabolic processes, oxidative phosphorylation or amino acid synthesis. **(A and B)** Each sgRNA is designated by the target gene name and whether it is the first or second designed sequence (i.e. a or b). Gray shaded heatmap describes the results from previous TnSeq experiments^6^ as presented in the mycobrowser database (https://mycobrowser.epfl.ch/) and whether it is essential (E), non-essential (NE) or whether it has a growth defect (D). Pink shaded heatmap describes the predicted strength of PAM scores ranked between 1-15 based on previous studies by Rock et al^15^. Blue shaded heatmap describes the level of growth inhibition at increasing ATc concentrations relative to a no ATc control well. The level of inhibition is the mean of at least four replicate experiments. Orange shaded heatmap describes the log_10_ reduction in CFU/ml for each sgRNA between day 0 and day 5 at increasing concentrations of ATc. The log_10_ reduction in CFU/ml of inhibition is the mean of at least four replicate experiments.

### Genes involved in amino acid synthesis are essential but generally bacteriostatic

Transcriptional inhibition of genes involved in amino acid biosynthesis generally inhibited bacterial growth (i.e. *hisG, aroK, trpD/A/B, ilvB1/D, thrA, metX, lysA*). These essential phenotypes are consistent with previous Tn-seq experiments (Figure 3C)^6^. The majority were bacteriostatic with only *aroK* being bactericidal. Although the sgRNA *metX*_a reduced bacterial viability at 33 ng/ml ATc (i.e. 1.02 ± 0.92 log_10_ reduction), this phenotype was not observed across other concentrations or sgRNAs (Figure 3C). Only a single *argJ* sgRNA was able to inhibit bacterial growth at the highest ATc concentration, whilst sgRNAs targeting *serB2* and *glnA* failed to inhibit bacterial growth (Figure 3C)^6^. Non-essentiality calls for *ald* and *lysE* are consistent with previous experiments (Figure 3C)^6^. In conclusion, the transcriptional inhibition of genes involved in amino acid synthesis generally inhibit bacterial growth, but have a bacteriostatic phenotype.

### Comparison of sgRNA activity identifies pathways of altered vulnerability

To identify cellular pathways more vulnerable to inhibition (i.e. requiring less ATc to inhibit bacterial growth) all sgRNAs were grouped according to biological function and the ATc MIC of each active sgRNA (i.e. that inhibited bacterial growth) was compared. To account for potential variation between experiments, the ATc MIC of sgRNAs was compared to the ATc MIC of the *mmpL3*_a sgRNA that was used as a positive knockdown control in all experiments (Figure 4B). The majority of sgRNAs targeting cell wall synthesis or core cellular process were as active as *mmpL3*_a (i.e. within a log2 change) (Figure 4A). Several sgRNAs targeting cell wall synthesis, core process and metabolic genes were more active (i.e. >-1 log2 change), representing pathways of potentially increased vulnerability (Figure 4A-B, red symbols). Interestingly, *eno*, which converts 2-phosphoglycerate to phosphoenolpyruvate as part of glycolysis, was the most vulnerable of all assessed sgRNAs (Figure 4A-B). A significant proportion of sgRNAs targeting metabolism, oxidative phosphorylation and amino acid synthesis genes were less active than *mmpL3*_a (i.e. >1 log2 change), representing biological pathways of potentially reduced vulnerability (Figure 4A-B, blue symbols). Dose response curves for five sgRNAs of either increased or reduced vulnerability highlight this shift in ATc MIC (Figure 4B).

**Figure 4:**
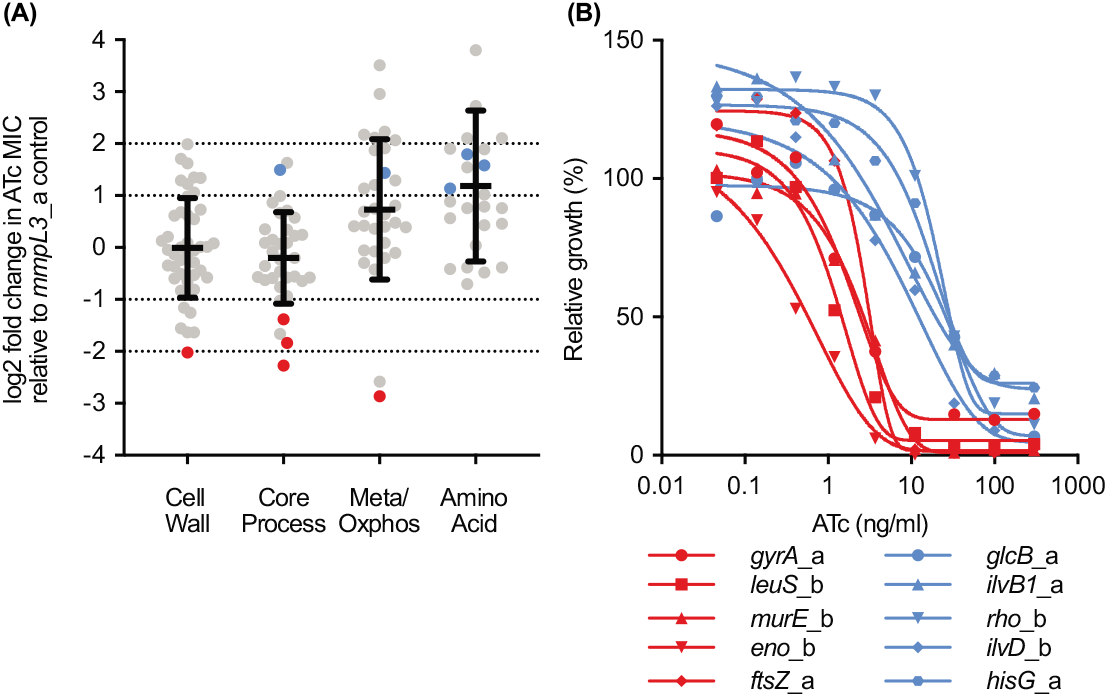
Comparison of target vulnerability across cellular pathways. (A) Plotted data is the average log2 fold change in ATc MIC compared to the *mmpL3*_a sgRNA for each sgRNA that results in at least a growth impairment (i.e. <50% relative to no ATc control) at the maximum ATc concentration. Data is the average fold change from at least two experiments that individually included biological duplicates (i.e. at least 4 replicates). sgRNAs are grouped according to their predicted or validated cellular function. Average sgRNA fold change values are presented in table S1. The average log2 fold change for a cellular pathway is represented by a solid black line. (B) Dose response curves for selected sgRNAs. Red and blue dots in A correspond to red and blue lines in B. Dose response curves are the average of at least 4 replicate experiments.

## Discussion

There remains an urgent global need for novel therapeutic agents to treat both drug susceptible and resistant strains of *M. tuberculosis*. Biological pathways that are essential, highly vulnerable to inhibition and have a bactericidal outcome are likely to have the greatest clinical impact. High-throughput and targeted genetic deletion strategies have been useful in predicting gene essentiality. However, each strategy is subject to individual limitations that include an inability to determine target vulnerability, differentiation between lethal and static outcomes or issues with scalability. By coupling CRISPRi to assessments of growth and viability this study overcomes these individual limitations to identify variations in target vulnerability and consequences on cellular viability for a diverse array of biological pathways.

There was a strong correlation between genetic essentiality and CRISPRi mediated growth inhibition when targeting genes involved in cell wall synthesis or core cellular processes. The majority of cell wall synthesis or core cellular encoding genes were equally vulnerable to transcriptional inhibition and bactericidal. Genes involved in mycolic acid synthesis, DNA metabolism, transcription, translation, tRNA synthesis, protein homeostasis or cell division had the strongest bactericidal phenotype (i.e. both level of reduction and levels of ATc needed to induce killing). Inhibition of peptidoglycan and arabinogalactan biosynthesis had differing effects on cell viability depending on the gene being targeted, with bactericidal phenotypes generally being weaker. Due to the polar effects of CRISPRi^15,16^ the repression of *murE* and *murX* is likely to repress multiple downstream genes within the *dcw* operon (including, *murD, murF, murG, ftsW* and *ftsQ*)^19^. Consequently, bactericidal phenotypes for *murE* and *murX* may not be reflective of single target gene repression. Similarly, the polar effects of CRISPRi suggest that transcriptional inhibition of both gyrase subunits, as would be facilitated by the *gyrB* sgRNA on the *gyrBA* operon but not *gyrA* sgRNA, is needed for lethality. Furthermore, both sgRNAs targeting *rho* have low PAM scores (i.e. 9 and 11), which may explain the bacteriostatic phenotype compared to previously published bactericidal *rho*-DUC mutants^20^. Combined, the results of this study support the continued development of novel drugs that target proteins involved in cell wall synthesis and core cellular processes.

Genes involved in metabolic processes, oxidative phosphorylation or amino acid synthesis, were on average less vulnerable to transcriptional inhibition. Genes that inhibited bacterial growth were generally bacteriostatic, with the bactericidal exceptions of *glcB* (i.e. malate synthase)^21^, *atpE/B* (i.e. ATP synthase operon)^16^, *aroK* (i.e. shikimate biosynthesis) and *ndh* (i.e. NDH2 dehydrogenase). Recent studies have demonstrated that the consequences of inhibiting individual metabolic genes when using CRISPRi can be mitigated by metabolic buffering, to limit global effects on bacterial metabolism^22^. Consequently, several cell divisions may be needed to deplete enzyme products below critical levels. Previous studies in *E. coli* demonstrated that metabolic buffering resulted in time dependent delays in inhibition of bacterial growth, with only a small proportion of metabolic genes having rapid inhibitory phenotypes^22^. We hypothesize that metabolic buffering in *M. tuberculosis* may explain why many predicted essential metabolic, oxidative phosphorylation or amino acid synthesis genes fail to inhibit bacterial growth or have a significantly reduced vulnerability when targeted by CRISPRi. Metabolic buffering may also explain why *metX, thrA* and *ilvB1* were bacteriostatic, yet in previous studies deletion mutants had bactericidal phenotypes ^2,23,24^. Consistent with this, previous observations of *metX* and *thrA* lethality are observed post 5-days, whilst our assay assessed viability at 5-days ^2,23^. Media choice also plays an important role in the observation of phenotypes associated with metabolic mutants ^21,25,26^. Consequently, the use of OADC (i.e. glucose and oleic acid as carbon sources) and exogenous pantothenic acid in our experiments to supplement *M. tuberculosis* mc^2^6230 (∆*panCD*) may alter or mitigate the phenotypes of some metabolic, oxidative phosphorylation or amino acid transcriptional knockdowns. Furthermore, the lack of phenotype for sgRNAs targeting *birA* is likely due to the presence of biotin in 7H9 media.

In our study, numerous previously defined non-essential genes had significant impacts on bacterial growth. In contrast to the metabolic buffering that may mitigated the phenotypes of essential genes, we hypothesize that a lack of metabolic buffering in some cases may delay the necessary metabolic remodelling that is required to facilitate growth in the absence of these non-essential genes. Consequently, previously constructed knockouts or Tn-seq mutants of of *icl1*^27^, *qcrB*^25^, *ctaC*^25^, *dlaT*^28^ and *ndh*^25^ may reflect an adapted metabolic state that in our study are defined as being essential for bacterial growth.

We acknowledge that variations in genetic vulnerability may be a result of variation in CRISPRi transcriptional repression. It is also possible that differences in bacterial phenotypes between sgRNAs (e.g. one static and one cidal) may be due to differences in sgRNA efficacy. Previous studies have highlighted the influence of PAM sequence, location of sgRNA within the genetic target and sgRNA sequence on dCas9 *Streptococcus pyogenes* activity^29-31^. Whilst we selected the strongest predicted PAM sequence and sgRNA location does not influence activity of the dCas9_Sth1_orthologue^15^, more work is required to determine the influence of sgRNA sequence on dCas9_Sth1_ activity. Off-target effects are also unlikely to be a confounding factor, as previous work as suggested limited off-targeting for the dCas9_Sth1_ orthologue^14^. Despite this, we speculate that variable levels of transcriptional repression are not a confounding factor on our overall conclusions as CRISPRi studies in *E. coli* and *B. subtilis* have demonstrated that genes involved in peptidoglycan synthesis and translation are more vulnerable to inhibition than metabolic genes^31^.

There is good correlation between the phenotypes of transcriptional inhibition and many known antibiotics. For example, the bactericidal phenotype of isoniazid, a cornerstone of current treatment regimens, correlates well with transcriptional inhibition of its target *inhA*^*32*^. Interestingly, transcriptional inhibition of *rpoB* produced a strong bactericidal phenotype, whilst rifampicin, a known inhibitor of RpoB, only achieves bacterial killing when in significant excess of the MIC^32^. Whilst it is possible that there are differences between the consequences of transcriptional and chemical inhibition, these results suggest that rifampicin may be a poor inhibitor of RNA polymerase *in vitro*. Combined, the results of our current study have provided novel insights into the vulnerability and lethality of inhibiting different biological processes in *M. tuberculosis*. These results will be a valuable resource for the mycobacterial research community that will help to advance both basic biology and the advancement of novel drug targets.

## Methods

### Bacterial strains and growth conditions

*Escherichia coli* MC1061 was grown in or on luria broth (LB) media or agar (1.5%) at 37°C and shaking at 200 rpm when required. *M. tuberculosis* strain mc^2^6230 (∆*panCD*, ∆RD1) ^33-36^, was grown and maintained in 7H9 liquid media or on 7H11 solid media supplemented with OADC (0.005% oleic acid, 0.5% bovine serum albumin, 0.2% dextrose, 0.085% catalase) and pantothenic acid (25 µg/ml) and incubated at 37°C. Liquid cultures were supplemented with 0.05% tyloxapol (Sigma) and grown with shaking at 140 rpm. *M. tuberculosis* strain mc^2^6230 is a BSL2 avirulent auxotroph that has been approved for use under BSL2 containment at the University of Otago. When necessary media was supplemented with Kanamycin at 50 µg/ml for *E. coil* and 25 µg/ml for *M. tuberculosis*. Anhydrotetracycline (ATc, Sigma) was solubilized in 70% ethanol and added to experiments at the stated concentrations.

### Construction and transformation of CRISPRi plasmids

A 20-25 bp sequence downstream of permissible PAM sequences targeting the non-template strand of target genes of interest were identified^15^. For each target gene, the two sgRNAs based on PAM score were manually selected. Target sequences were ordered as oligos with GGGA and AAAC overhangs respectively (Table S1), and cloned into pJLR965 using BsmB1 and golden gate cloning as previously described^13^. Plasmids were cloned into *E. coli* and validated with sanger sequencing. Confirmed CRISPRi plasmids were electroporated into *M. tuberculosis* strain mc^2^6230 following previously established protocols^13,16^.

### CRISPRi phenotypic assessment of essentiality and viability

To determine the consequences of targeted gene repression on bacterial growth, phenotypic assays were performed as follows (Figure 1). *M. tuberculosis* mc^2^6230 strains containing CRISPRi plasmids were grown and maintained in 7H9-supplemented media with KAN. Cultures were diluted initially to an OD_600_of 0.1 in 7H9-supplemented media (+KAN) in a deep well 96 well plate. Cultures were diluted again to a final OD_600_ of 0.01 in a deep well 96 well plate. 96 well assay plates were prepared with a 3-fold dilution of ATc along the X-axis starting at 300 ng/ml of ATc in column 2 with a starting inoculum of OD_600_0.005 (Figure 1). This was achieved by adding 50 µl of 7H9-supplemented media (+KAN) to all wells of columns 3-11 except row A-H. 75 µl of 7H9-supplemented media (+KAN) containing the starting concentration of ATc (i.e. 600 ng/ml ATc) was added to column 2 except row A-H. ATc was diluted along the horizontal axis of the 96 well plate, transferring 25 µl between columns, down to column 10. Column 11 was used as a no ATc control. Columns 1 and 12 and rows A-H contained 100 µl of 7H9-supplemented media (+KAN) as contamination and background controls. Fifty µl of culture adjusted to an OD_600_ 0.01 was added to wells 2-11 of individual rows of a 96 well plate to achieve a starting OD_600_ of 0.005. Each row represents the ATc dilution gradient for a single *M. tuberculosis* mc^2^6230 strain expressing a unique CRISPRi plasmids. All experiments included a non-targeting sgRNA (i.e. pJR965) and a *mmpL3_*a targeting sgRNA as a negative and positive essential-bactericidal control^13^.

To assess gene essentially, duplicate plates were grown at 37°C without shaking for 7-days. OD_600_ was measured using a Varioskan-LUX microplate reader. The minimal inhibitory concentration (MIC) of ATc was determined using OD_600_ reads from duplicate plates relative to the growth of the no-ATc control, using a non-linear fitting of data to the Gompertz equation^37^. Targeted genes were defined as either essential (i.e. <25% growth relative to no ATc control), growth impairing (i.e. <50% growth relative to no ATc control) or non-essential (i.e. >50% growth relative to no ATc control). In this study, the vulnerability of genes to transcriptional inhibition is defined by the ATc MIC, i.e. the lower the MIC the more vulnerable that gene is to CRISPRi mediated transcriptional inhibition.

Duplicate 96 well assay plates were also set up to determine the consequences of targeted gene repression on bacterial viability by taking colony forming units on day 0 and day 5. Previous work has demonstrated that day 5 allows for the detection of bacterial killing prior to the emergence of non-responsive CRISPRi mutants^13,16^. Briefly, at day 0 hrs a 4-point ten-fold dilution of the 0.01 diluted culture was performed in 7H9 base media (i.e. not supplemented), with 5 µl of each dilution spotted onto to 7H11-supplemented (+KAN) agar plates. At Day 5, approximately 100 µl of culture was removed from columns 2-5 of each row and transferred to a new 96 well plate to be diluted. A 4-point ten-fold dilution gradient was performed in 7H9 base media (i.e. not supplemented), and 5 µl of each dilution was spotted onto to 7H11-supplemented (+KAN) agar plates. Plates were incubated at 37°C for 4-5 weeks, at which point colonies were counted. Essential genes with bacteriostatic consequences resulted in no change in CFU/ml relative to 0 hrs, whilst bactericidal consequences produced at least a 1 log reduction in CFU/ml relative to the day 0 inoculum.

## Acknowledgments

This research was financially supported by the Maurice Wilkins Centre for Molecular Biodiscovery, the Marsden Fund (Royal Society of New Zealand) (grant number UOO1807). and the Health Research Council of New Zealand (grant number 20/459).

We have no conflicts of interest to declare.

